# Metagenomics and qPCR analysis of aerobic metabolic TCE degrading bacteria

**DOI:** 10.1101/2025.07.31.667672

**Authors:** Steffen Hertle, Lara Stelmaszyk, Annika Behrendt, Johannes Ho, Xiaojun Zhang, Zhao He-Ping, Timothy M. Vogel, Andreas Tiehm, Azariel Ruiz-Valencia

## Abstract

Chloroethenes are common groundwater contaminants. While they were initially considered non-biodegradable, different pathways for both, anaerobic and aerobic degradation have been discovered over the last 40 years. Anaerobic reductive dechlorination of tetrachloroethene and trichloroethene (TCE) leads to the formation and possible accumulation of the toxic intermediate products, dichloroethene (DCE) and vinyl chloride (VC). In contrast, aerobic metabolic processes result in mineralization of the contaminants without the formation of stable intermediates. Therefore, productive aerobic degradation processes can have a considerable advantage, depending on site conditions and chloroethene molecule. The potential for aerobic degradation processes can be assessed by molecular biological analysis. In our study, aerobic metabolic TCE enrichment cultures, bioaugmented groundwater, and a microcosm with intrinsic aerobic TCE degradation potential were analyzed. Amplicon sequencing and shotgun metagenomics were used to identify the *Rhodocyclaceae* bacterial family in all samples, as well as functional gene sequences, which were relatives of monooxygenases and dehalogenases. Based on these sequencing data, qPCR detection methods for TCE assimilating *Rhodocyclaceae* (*Rho*) as well as for different functional genes that were associated with monooxygenases (*moABC*) and haloacid dehalogenases like hydrolases (*hdlh*) have been established. The specificity of the qPCR methods was demonstrated with a variety of environmental samples and different chloroethene degrading bacterial cultures. The new metagenomic insights and qPCR methods facilitate the assessment of biodegradation potential and monitoring of aerobic metabolic TCE degradation at contaminated sites.

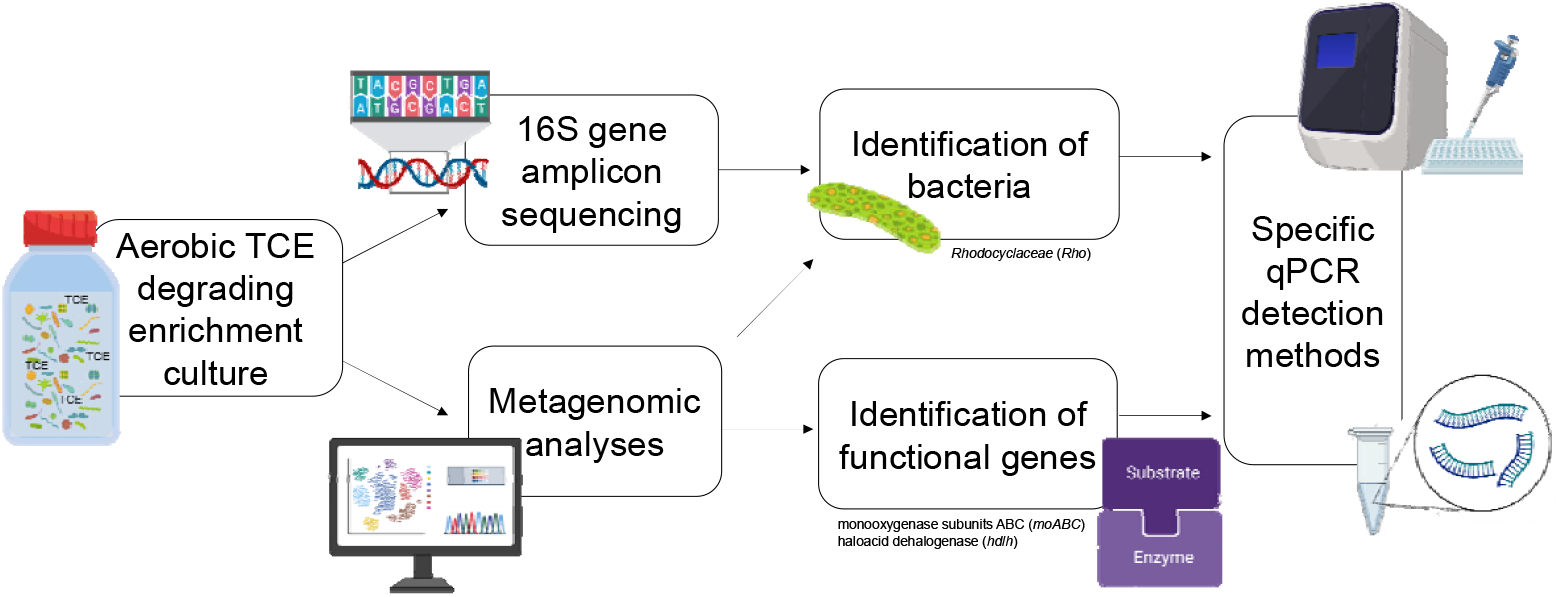

**Highlights:**

- *Rhodocyclaceae* bacteria were identified in all aerobic TCE degrading samples
- Functional monooxygenase and dehalogenase like genes were detected
- qPCR methods for monitoring of aerobic metabolic TCE degradation were developed
- Specificity of qPCR methods was shown with different aerobic chloroethene degrading cultures and environmental samples

## 1 Introduction

Due to the use of chloroethenes in different industries like dye-production, dry-cleaning, metal works etc., they have become common groundwater contaminants worldwide. Used for their beneficial properties as a solvent, their recalcitrance and ecotoxicity has curtailed their use, although contaminated sites remain. While initially considered non-biodegradable, different biodegradation pathways have been identified starting in the 1980s. The higher-chlorinated ethenes, tetrachloroethene (perchloroethene, PCE) and trichloroethene (TCE) were shown to be degraded microbially during anaerobic reductive dechlorination (Vogel *et al*. 1985). In the reductive dechlorination pathway, the chloroethenes are transformed through a cascade of dechlorination steps, where a chlorine-substituent is replaced by hydrogen-atom (Freedman and Gossett 1989). In the case of PCE, the complete reductive dechlorination results in the temporary presence of the lower chlorinated ethenes TCE, cis-1,2-dichloroethene (cDCE) and vinylchloride (VC) (Bradley and Chapelle 2007) before being completely dechlorinated to ethene in some cases. Due to inadequate environmental conditions as well as lack of competent bacteria at contaminated sites, the reductive dechlorination can stall, resulting in the accumulation of the toxic intermediate products cDCE and VC in the subsurface (discussed in Tiehm and Schmidt, 2011). Additionally, sulfate reduction and methanogenesis have to be considered if excess auxiliary substrates are applied at contaminated sites to stimulate reductive dechlorination (Tiehm and Schmidt 2011).

In addition to the reductive dechlorination pathway, aerobic processes can be utilized to degrade the chloroethenes TCE, cDCE and VC. The aerobic pathways can further be differentiated into co-metabolic and metabolic pathways. Aerobic co-metabolic pathways require an auxiliary substrate (*e*.*g*. methane (Hazen *et al*. 2009), ammonium (Vanelli *et al*. 1990), propane (Arp *et al*. 2001), VC (Verce *et al*. 2002)), as growth substrate. The co-metabolic chloroethene degradation is fortuitous (Mattes *et al*. 2010) and with low efficiency as transformation yields for co-metabolic degradation of TCE range from 0.003-0.068 mg_TCE_/mg_Aux_ depending on the auxiliary substrate (Chang and Alvarez-Cohen 1995; Tovanabootr and Semprini 2010). VC (Hartmans and Bont 1992), cDCE (Jennings *et al*. 2009) and TCE (Schmidt *et al*. 2014) can be utilized in aerobic metabolic processes as growth substrate as well as an energy source, resulting in mineralization of the contaminants without stable and undesirable intermediate products. Therefore, this process has clear advantages over both, reductive dechlorination and aerobic co-metabolic degradation.

In order to demonstrate field site potential for the different degradation steps, molecular-biological analysis have been established as a line of evidence for reductive dechlorination and specific monitoring (Kranzioch *et al*. 2015; Michalsen *et al*. 2022). By applying qPCR-methods, proof of the presence of different bacteria and the different functional genes encoding for the enzymes directly involved in reductive dechlorination steps can be provided rapidly. Compared to the reductive dechlorination pathway, aerobic metabolic pathways are less well studied. Since different bacteria and functional genes have been identified for VC-(Jin and Mattes 2010) and cDCE-(Giddings *et al*. 2010) degradation, bacteria capable of aerobic TCE degradation are still unknown.

A mixed bacterial culture from the SF-site, where the aerobic metabolic TCE-degradation was first observed (Schmidt *et al*. 2014), was successfully used in bioaugmentation experiments (Gaza *et al*. 2019). In addition, the intrinsic aerobic metabolic TCE degradation has been observed at other sites (Willmann *et al*. 2023). The qPCR-detection is a useful tool in the monitoring and evaluation of bioaugmentation technologies. Using qPCR, more potential sites can be identified for the aerobic metabolic degradation of TCE. To determine the possible qPCR targets, amplicon sequencing and metagenomic approaches were applied to a range of samples with observed aerobic metabolic TCE degradation.

The objectives of the study were

1. to identify bacteria and functional genes involved in the aerobic metabolic degradation of TCE; and
2. to establish qPCR methods based on sequencing data, creating a method to detect and monitor aerobic TCE degrading bacteria and relevant functional genes.

## 2 Material and Methods

### 2.1 Incubation details

All samples used here came from sealed bottle experiments, where TCE was the only source of carbon and aerobic conditions were maintained with air filled headspace. The bottles were stored in the dark at room temperature. The different conditions included enrichment cultures originating from the SF-culture that was maintained in mineral medium (MM) with TCE as sole carbon source (Schmidt *et al*. 2014) grown over 14 years with around 50 transfers to fresh MM. In addition, two groundwater samples (one with added enrichment culture and one without) were included (Willmann *et al*. 2023). The different sealed bottles contained two batch cultures (SF-1a and SF-1), samples from column circulation systems with immobilized (SF-2a) and suspended biomass (SF-2, SF-3, SF-4), and groundwater with (SF-GW) or without added SF-culture (MC-GW). The degradation of the TCE and the production of chloride in the different bottles is shown in Figure S1.

The samples (from two time points) used for taxonomic analyses by amplicon sequencing came from a SF-culture in MM (SF-1a), SF-culture immobilized on quartz sand in MM recirculating in a column system (SF-2a), and groundwater from a TCE contaminated site (Willmann *et al*. 2023), bioaugmented with SF-culture (SF-GW). The samples used for metagenomic sequencing consisted of a different SF-culture in MM (SF-1), a SF-culture from a column outflow operated with MM (SF-2), a subsample of the outflow of a column with SF-culture enrichment immobilized on quartz sand (SF-3), a SF-culture enrichment immobilized on quartz sand (SF-4), and a microcosm with groundwater from a TCE contaminated site (Willmann *et al*. 2023) (MC-GW).

For the validation of the developed qPCR methods, two batch experiments were carried out with enrichment culture in MM and 10 mg/L TCE. qPCR samples were analyzed at three points (day 0, day 21 and day 36).

### 2.2 DNA Extraction

Fifty mL of the batch samples were concentrated by filtration through a polyethersulfonate membrane (Pall, Life Sciences Supor-200, 0.2 µm pore size, 45 mm diameter, Pall Corporation, Washington, USA). Filters were cut in half and stored at -20°C until DNA-extraction. DNA was extracted using the Fast DNA Spin Kit for Soil (MPBio, Santa Ana, USA) according to manufacturer’s instructions, except for the following adjustments: In step 4, 700 µL were discarded instead of 500 µL. In step 5, 700 µL were transferred into the spin filter tubes twice. For elution, 100 µL of a 56°C warm DES elution solution was used. The extracted DNA was stored at -20°C until measurement.

### 2.3 Amplicon sequence analysis

The workflow of amplicon sequence analysis is shown in Figure S2 (left). In short, the V4 region from the gene *rrs* that codes for the 16S rRNA gene was amplified using the Illumina primers ILL_515Fmod and ILL_806Rmod. The Illumina sequencing was performed by at ZIEL – Institute for Food and Health (TU Munich). The adapters for the barcoding were removed and reads that were too short (less than 208 bp) or too long (greater than 245 bp) were also removed by using cutadapt 4.4 (Martin 2011). The sequences were then analyzed following the DADA2 pipeline using the DADA2 1.28.0 (Callahan *et al*. 2016) package in R. The quality trimming was used to remove low-quality primer sequences, bases, and low-quality reads. Bimeric reads were identified in “consensus” mode and removed from forward reads. The sequences were annotated against the SILVA NR99 v138.1 database. Raw data from sequencing samples have been deposited in the European Nucleotide Archive (ENA) under accession numbers from ERS24548512 to ERS24548517 under the study PRJEB89694. This workflow was applied to samples SF-1a, SF-2a, SF-GW.

### 2.4 Shotgun metagenome analysis

Metagenomic sequencing analyses were carried out in triplicates for each sample (SF-1, SF-2, SF-3, SF-4, MC-GW; Figure S2, right). Extracted DNA was sheared to□∼300 bp using the Nextera XT kit and libraries were constructed using Nextera XT v2 Index Kit A. The products were purified by using the AMpure XP beads and quantified afterwards on a μDrop Plate reader (Thermo Scientific). The size-selection was performed with the AMpure XP beads, and the final library pool was quantified with the Bioanalyzer High Sensitivity DNA chip in an Agilent 2100 Bioanalyzer. The samples were sequenced in triplicate using a Mi-Seq sequencer (Illumina, USA) according to the protocol given by the manufacturer.

The sequences obtained were filtered to remove sequence noise (Minoche *et al*. 2011) and co-assembled using MEGAHIT v1.2.9 in sensitive mode with a minimum contig length of 1000 bp (Li *et al*. 2015). Bowtie2 was used to map the contigs and create the sample profile of each sample. The metagenomic workflow for ANVI’O was then used to visualize and manually bin the contigs. The binning was performed based on their sequence composition and differential coverage. The taxonomy of the bins was estimated by using the single-copy core gene sequences and comparing them with the GTDB database. Finally, the functions were annotated using KEGG and PFAM databases. Raw sequencing data from the different samples has been deposited in the European Nucleotide Archive (ENA) under accession numbers from ERS20167400 to ERS20167426 and under the study PRJEB76220.

### 2.5 Primer Design

The design of the primer *Rho* for qPCR analyses was based on the amplicon sequencing. The bioinformatic analysis to obtain the operational taxonomic units (OTUs) for the primer design of the predominant TCE degrading bacterial families was conducted using the method of Reitmeier *et al*. (2020). The design of the primers for the functional genes was based on metagenomic analysis. The primers were designed using Primer-Blast (NCBI) with the product size of max. 300 bp, annealing temperature between 57-63°C with an optimum at 60°C with a custom based fasta file including the target sequence and the specificity check limited to bacteria. The specific annealing temperatures of the primers were determined experimentally by quantitative real-time PCR with temperature gradient from 55°C to 70°C.

### 2.6 Quantitative real-time PCR

Quantitative real-time PCR analysis (qPCR) was performed with Rotor-Gene Q (Qiagen, Hilden, Germany) cycler. The 10 µL reaction consisted of 5 µL SsoAdvanced Universal SYBR Green Supermix (BioRad, Hercules, USA), 3.2 µL nuclease-free water, 0.4 µL of the 10 nM forward and reverse primer and 1 µL of the DNA-template. Initial denaturation was held at 98°C for 120 s following 45 cycles of (1) denaturation (98°C, 15 s), (2) annealing (primer specific temperature according to Table 1, 15 s) and (3) elongation (72°C, 20 s). All samples and standards were analyzed in duplicates. The qPCR results were analyzed using the Rotor Gene Q Series Software. For quality control, R^2^ of the standard curve as well as the amplification efficiency were determined and melt curve analysis was performed. Only qPCR experiments with R^2^ values >0.99 and efficiencies between 90 % and 105 % were considered. Amplification products were verified via QIAxcel® Advanced system (Qiagen). The limit of quantification (LOQ) was 2 gene copies/mL

**Table 1.**
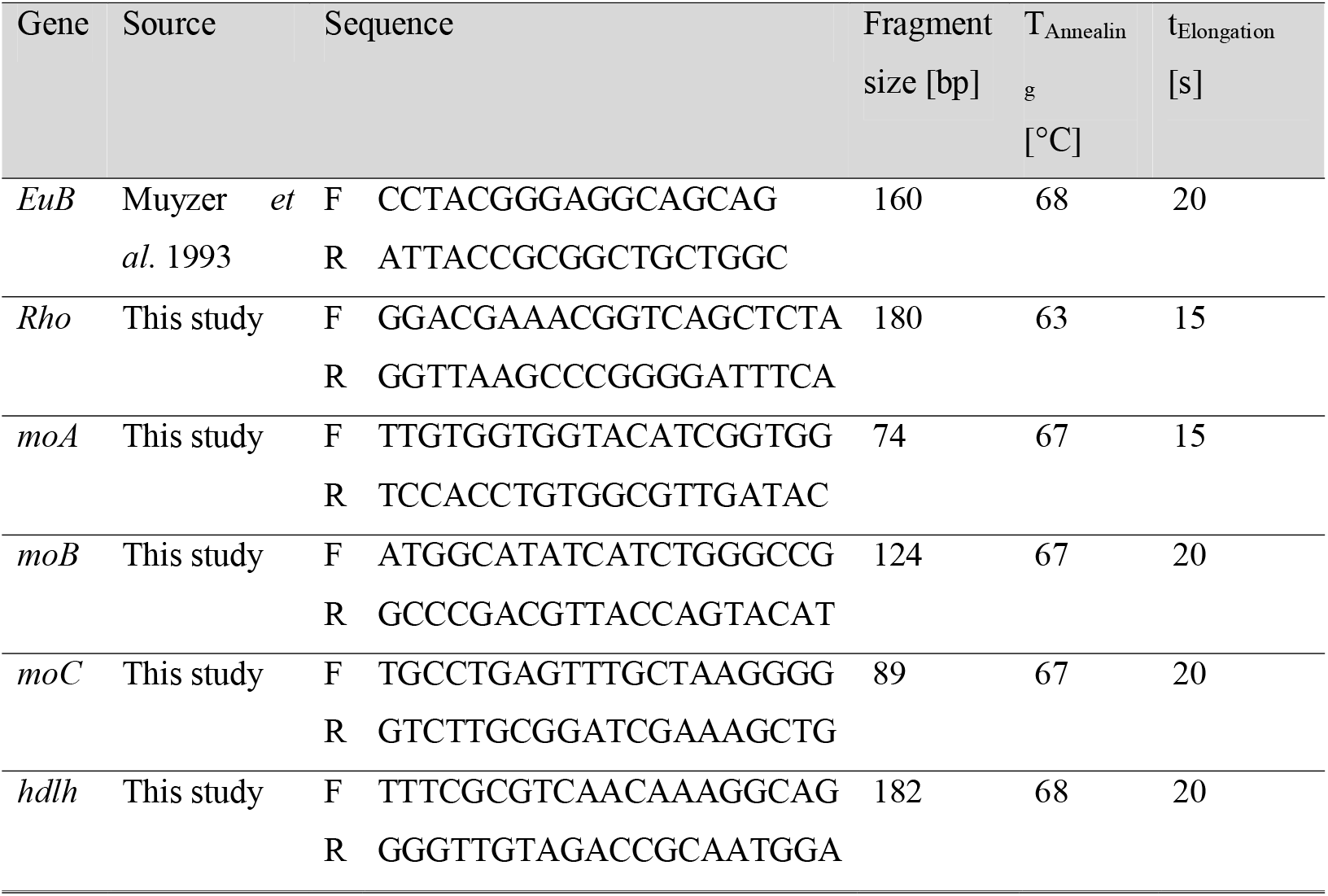
Primers and qPCR conditions for evaluating the presence of Rhodocyclaceae and putative TCE aerobic metabolic degradation genes. Sequences of the primers are given in 5’-3’-notation.

**Table 2.** Gene copy numbers detected in aerobic chloroethene enrichment cultures and environmental samples. Background information and groundwater analyses of the sites and batches are given in Table S1.

| Sample | gene copies [1/mL] |  |  |  |  |  |
| --- | --- | --- | --- | --- | --- | --- |
|  | *EuB* | *Rho* | *moA* | *moB* | *moC* | *hdlh* |
| Aerobic VC degrading culture (1) | 8.3e+6 | <LOQ | <LOQ | <LOQ | <LOQ | <LOQ |
| Aerobic VC degrading culture (2) | 3.4e+5 | <LOQ | <LOQ | <LOQ | <LOQ | <LOQ |
| Aerobic cDCE degrading culture (1) | 2.9e+4 | 3.0e+1 | <LOQ | <LOQ | <LOQ | <LOQ |
| Aerobic cDCE degrading culture (2) | 1.4e+6 | 9.5e+1 | <LOQ | <LOQ | <LOQ | <LOQ |
| Aerobic TCE degrading culture | 1.1e+7 | 4.3e+4 | 1.7e+4 | 1.6e+3 | 1.3e+4 | 1.7e+4 |
| Active TCE degrading site (Sample 1) | 2.7e+5 | 7.2e+4 | 1.1e+4 | 5.9e+2 | 2.7e+3 | 6.4e+3 |
| Active TCE degrading site (Sample 2) | 2.1e+5 | 7.2e+4 | 2.7e+2 | 3.3e+1 | 1.9e+2 | 1.7e+2 |
| Municipal waste water | 3.0e+6 | 6.0e+3 | <LOQ | <LOQ | <LOQ | <LOQ |
| BTEX/ PAH/ NSO-HET contaminated site 1 | 4.4e+4 | 1.5e+2 | <LOQ | <LOQ | <LOQ | <LOQ |
| BTEX/ PAH/ NSO-HET contaminated site 2 (Sample 1) | 6.1e+4 | 2.2e+2 | <LOQ | <LOQ | <LOQ | <LOQ |
| BTEX/ PAH/ NSO-HET contaminated site 2 (Sample 2) | 6.4e+5 | <LOQ | <LOQ | <LOQ | <LOQ | <LOQ |
| BTEX/ PAH/ NSO-HET contaminated site 3 (Sample 1) | 4.4e+7 | 6.1e+3 | <LOQ | <LOQ | <LOQ | <LOQ |
| BTEX/ PAH/ NSO-HET contaminated site 3 (Sample 2) | 2.4e+5 | 1.2e+3 | <LOQ | <LOQ | <LOQ | <LOQ |
| BTEX/ PAH/ NSO-HET contaminated site 3 (Sample 3) | 1.0e+4 | 4.8e+2 | <LOQ | <LOQ | <LOQ | <LOQ |

VC - Vinylchloride, cDCE = cis-1,2-dichloroethene LOQ = Limit of Quantification

### 2.7 Chemical Analysis

The batch experiments were conducted in mineral salt medium (MSM) based on the method described by Schmidt *et al*. (2014) with TCE (99,6 % purity, Sigma Aldrich) as sole source of carbon. TCE was measured with an Agilent Technologies (Waldbronn, Germany) 7890A GC system equipped with electron capture detector (ECD), flame ionization detector (FID) and 7697A headspace sampler. Values represent the average of duplicate measurements. For quality control, external standards have been analyzed with each run. Recovery rate of the standards was between 95 and 105 %. The limit of quantification for TCE was 0.7 μg/L; the limit of detection was 0.2 μg/L. The chloride concentration was determined with a Metrohm 761 compact ion chromatograph (Filderstadt, Germany) equipped with a conductivity detector and a MetrosepA-Supp-5 column with a limit of quantification of 1 mg/L and a limit of detection of 0.3 mg/L. Standard error of the analysis was 5%. Sample analysis were accompanied by the analysis of external standards for quality control. Recovery rate of the standards was between 95-105 %.

## 3 Results and Discussion

### 3.1 Identification of putative TCE degrading *Rhodocyclaceae* family by 16SrRNA gene amplicon analyses and metagenomic approaches

Bioinformatic analysis of the 16SrRNA gene amplicon from the three bottles, enrichment cultures (SF-1a, SF-2a); and a SF-culture-augmented groundwater batch (SF-GW) resulted in allocated sequences to several different bacterial families. The abundance of the most prevalent bacterial family belonged to *Rhodocyclaceae*. The *Rhodocyclaceae* family abundance increased from 12.5 % (SF-1a) and 5.1 % (SF-2a) to 67.9 % (SF-1a) and 43.6 % (SF-2a), respectively, after inoculation and 71 days of TCE degradation in mineral medium. This family abundance increased from 0.3 % (SF-GW) to 19.8% (SF-GW) after bioaugmentation of groundwater and 117 days of incubation (Fig. 1).

**Figure 1.**
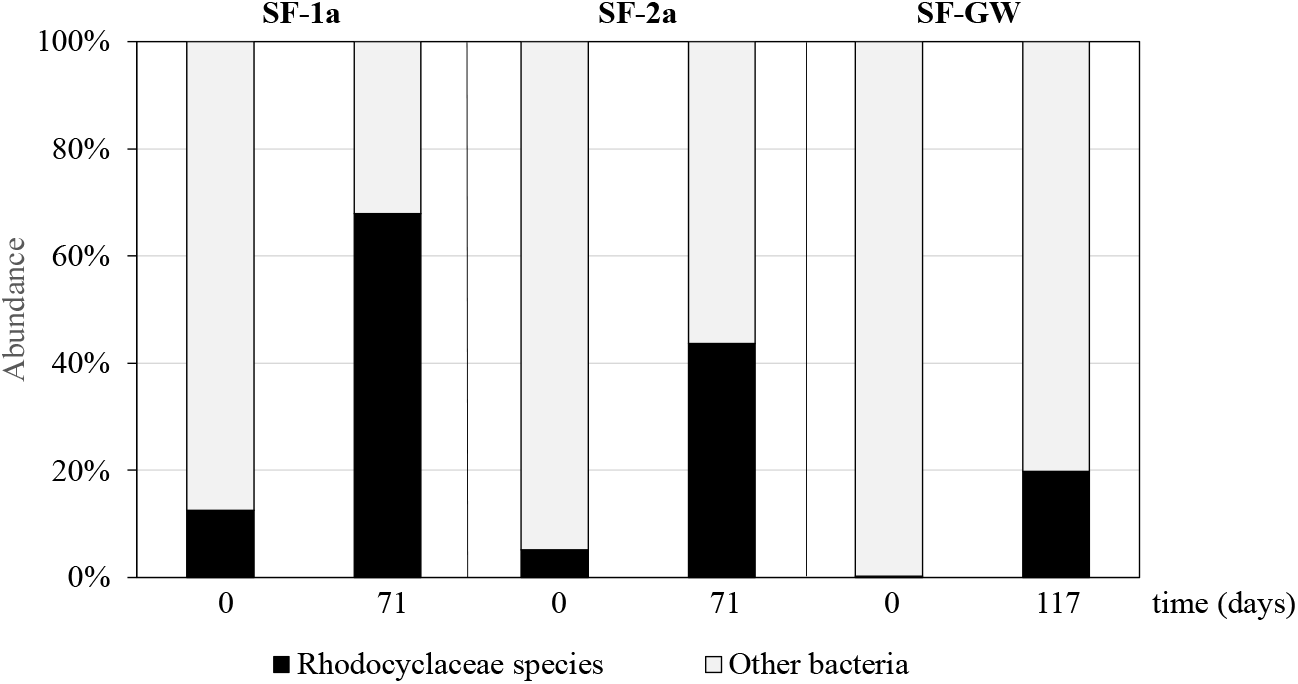
Shift of relative abundance of the Rhodocyclaceae species in batch experiments inoculated with the SF-culture and amended with TCE (SF-1a, SF-2a and SF-GW).

The metagenomic sequencing analysis (including assembly) of another five samples, described in section 2.1 (SF-1, SF-2, SF-3, SF-4, MC-GW), was consistent with these findings (Figure 2). The sequence analysis and assembly produced high-quality bins with > 80% completeness and < 8% redundancy of certain genomes. There was low variation of the taxonomic annotation of the single copy genes. The only bin with contigs presented in all five samples was annotated to the *Rhodocyclaceae* family, belonging to the order of *Burkholderiales*. Even with the high completeness of the bins, the genus could not be determined.

**Figure 2.**
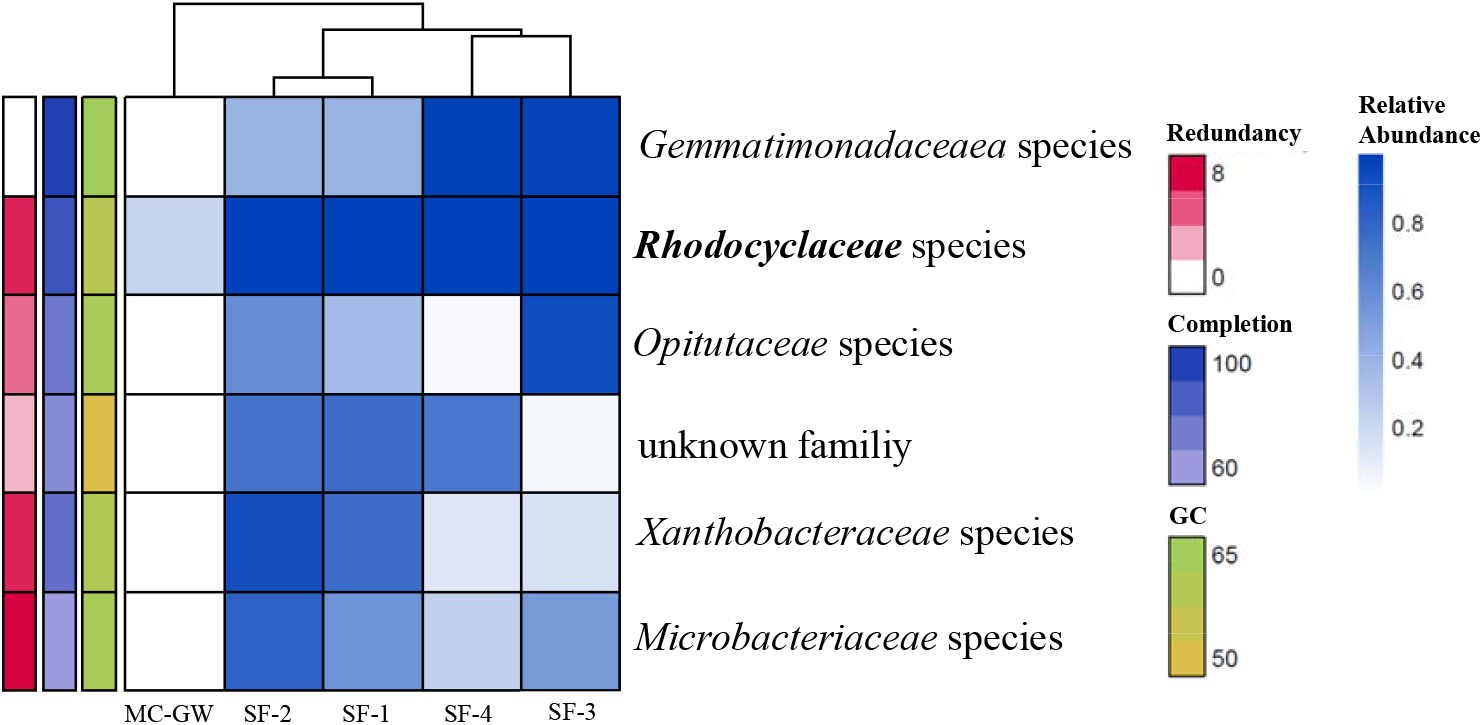
Relative abundance (blue shading for 0-15 normalized counts) and taxonomy for six bins assembled from all the samples with TCE showing that the bin annotated as Rhodocyclaceae was the most abundant family in all sequenced samples. Dendrogram organizes the samples according to the relative abundance of the bins for each sample (Euclidean distance and Ward linkage). Completion (60-100%), redundancy (0-8%) and GC content (50%-65%) are also shown.

### 3.2 Metagenomic analysis of functional genes involved in aerobic TCE degradation pathways

The metagenomic bins were analyzed for different genes that hypothetically could be involved in TCE degradation and in particular dehalogenases and monooxygenases. The sequences annotated for monooxygenases and dehalogenases in each bin/genome were used to identify potential TCE-degrading enzymes. A total of thirty-two (mono- and di-) oxygenase and two dehalogenase related enzymes were identified within the different bins (Figure S3). If the species within the *Rhodocyclaceae* family are responsible for the TCE degradation, the necessary genes should be present within its bin/genome. There were some dehalogenases and monooxygenases that were identified only within the *Rhodocyclaceae* bin, namely haloacetate dehalogenase (EC 3.8.1.3), methane/ammonia monooxygenase subunits A, B and C (EC 1.14.18.3) and choline monooxygenase (EC 1.14.15.7). Since the three subunits (A, B and C) for the particulate methane/ammonia monooxygenase (*amo*CAB/*mmo*CAB) operon (Figure) were identified in the *Rhodocyclacea* binned sequences and purified soluble methane monooxygenase (sMMO) was reported to degrade trichloroethene co-metabolically under aerobic conditions (Fox *et al*. 1990), these subunits could be involved in the aerobic metabolic degradation of TCE.

The annotated sequence found within the *Rhodocyclaceae* related sequences codes for a membrane bound particulate methane monooxygenase (pMMO) and, studies for TCE oxidation with pMMO extracted from methanotrophic bacteria and whole cells, describe an epoxide intermediate (Lontoh *et al*. 2000).

Two different sequences, corresponding to the subunit B of the translated protein, where the active site is located within the soluble region (Culpepper and Rosenzweig 2012), were annotated within the *Rhodocyclacea* bin. These gene and protein sequences annotated in the NCBI Reference Sequence Database were compared in a phylogenetic tree to determine how the monooxygenase subunit B sequences detected within *Rhodocyclaceae* family sequences are related to other subunit B sequences from other genera (Figure 3). The sequence present in the operon (split 33583) of the *Rhodocyclacea* bin did not cluster with any other reference sequence. One hypothesis for the lack of significant similarity could be that the monooxygenase gained the ability to catalyse the oxidation of TCE more efficiently than the monooxygenase evolved for oxidizing methane and thus, separated from the database enzymes through mutation. In addition, the second *pmo*B gene in the *Rhodocyclaceae* bin (split 33561) clustered near to ammonia oxidizing proteins. The length of this branch also indicates a significantly dissimilar sequence compared to the surrounding ammonia monooxygenases. The prolonged exposure to TCE may have changed the specificity of these enzymes involved in TCE degradation.

**Figure 3.**
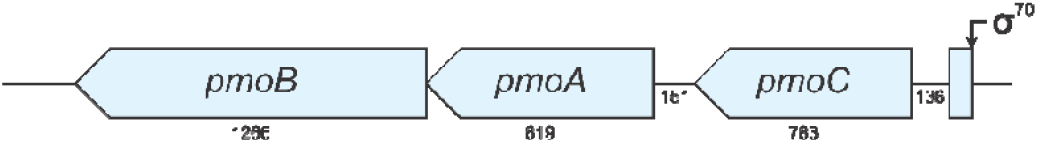
pmoCAB operon (with the subunit sizes in bp) for monooxygenase found in the Rhodocyclaceae bin.

**Figure 3.**
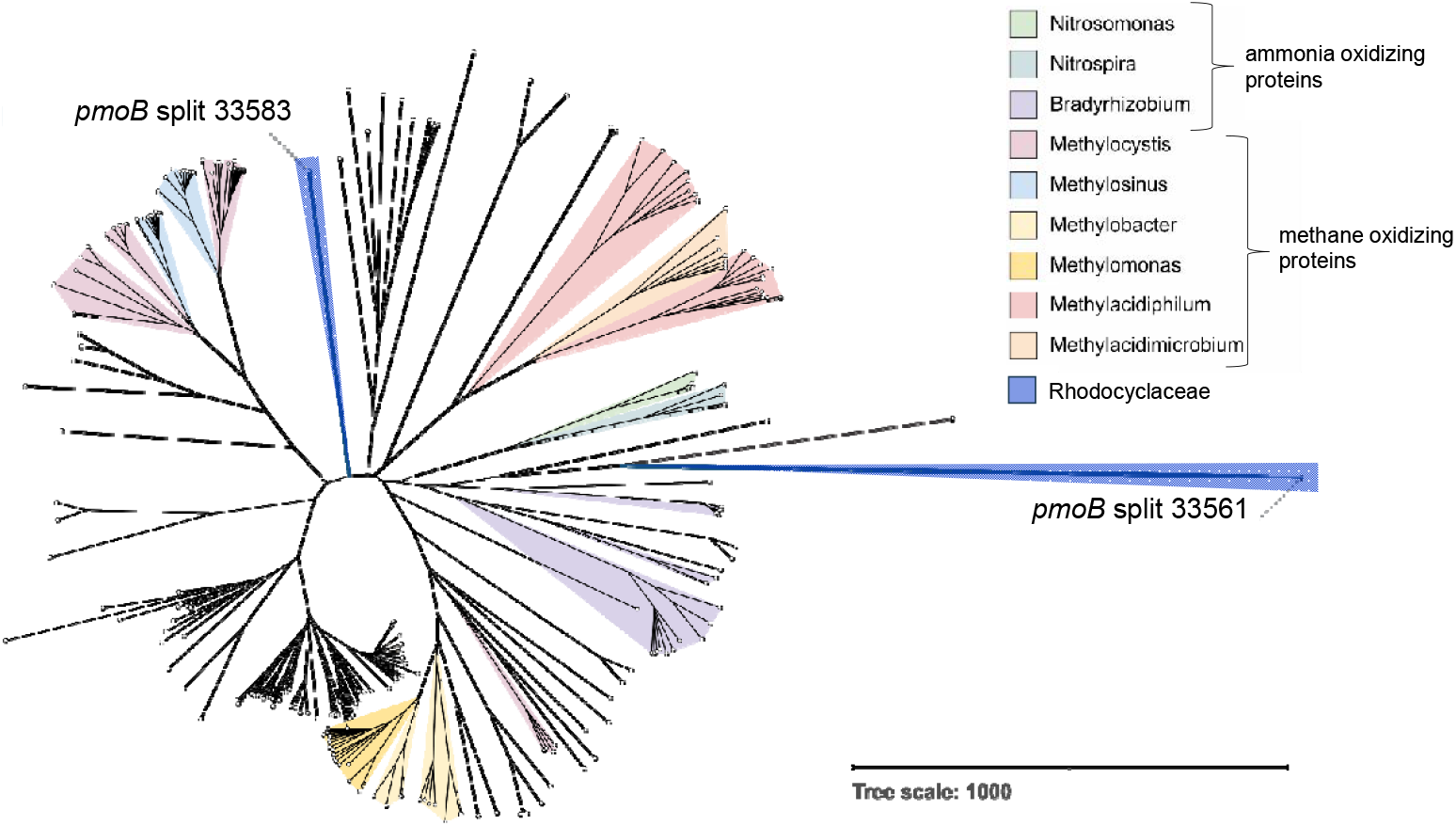
Unrooted neighbor joining phylogenetic tree from MuscleWS alignment for the translated amino acid sequences of pmoB from the Rhodocyclaceae bin and related sequences from the Reference Sequence database in NCBI.

A dehalogenase related enzyme might also be involved in the aerobic mineralization of TCE, since chloride is the main product of the TCE degradation (Schmidt *et al*. 2014). Two sequences coding for the haloacetate dehalogenase were found in the *Rhodocyclaceae* bin. Haloacetate dehalogenases are reported to catalyze the dechlorination of mono- and dichloroacetate (Weightman *et al*. 1985). Due to these identified enzyme encoding sequences in the *Rhodocyclaceae* bin dominating the TCE degrading enrichment cultures and groundwater batches, the sequences for monooxygenase subunit A, B and C (*moABC*), as well as the haloacetate dehalogenase (*hdlh*) were selected (shown in SI section 2) for establishment of a specific qPCR detection method.

### 3.3 Primer establishment and qPCR analysis in enrichment cultures

Based on the sequencing results, qPCR analyses were established for the 16S rRNA gene of the *Rhodocyclaceae* (*Rho)* and for the functional genes *moABC* (monooxygenases) and *hdlh* (dehalogenase) (Table 1). Additionally, “universal” bacterial 16S rRNA gene (*EuB*) was analyzed as a surragote parameter for overall bacterial biomass (Table 1).

A batch experiment was performed for 70 days to investigate the co-occurrence of metabolic aerobic TCE degradation and the quantity of genes (derived from the *Rhodocyclaceae* bin) implicated in the TCE degradation. As already reported previously (Schmidt *et al*. 2014, Gaza *et al*. 2019), the aerobic TCE degradation corresponds to the increase in chloride concentration (Figure 5, top). The specific newly established qPCR methods monitored the related growth of *Rhodocyclaceae* through increase of bacterial DNA and functional genes (Figure 4, bottom). Gene copy numbers reach a concentration of ∼10^5^ gene-copies/mL for the overall bacterial marker *EuB* and 10^4^ gene-copies/mL for the *Rhodocyclaceae* biomarker gene (*Rho*) and for putative TCE degradation genes as described above (*moA, moB, moC* and *hdlh*).

**Figure 4.**
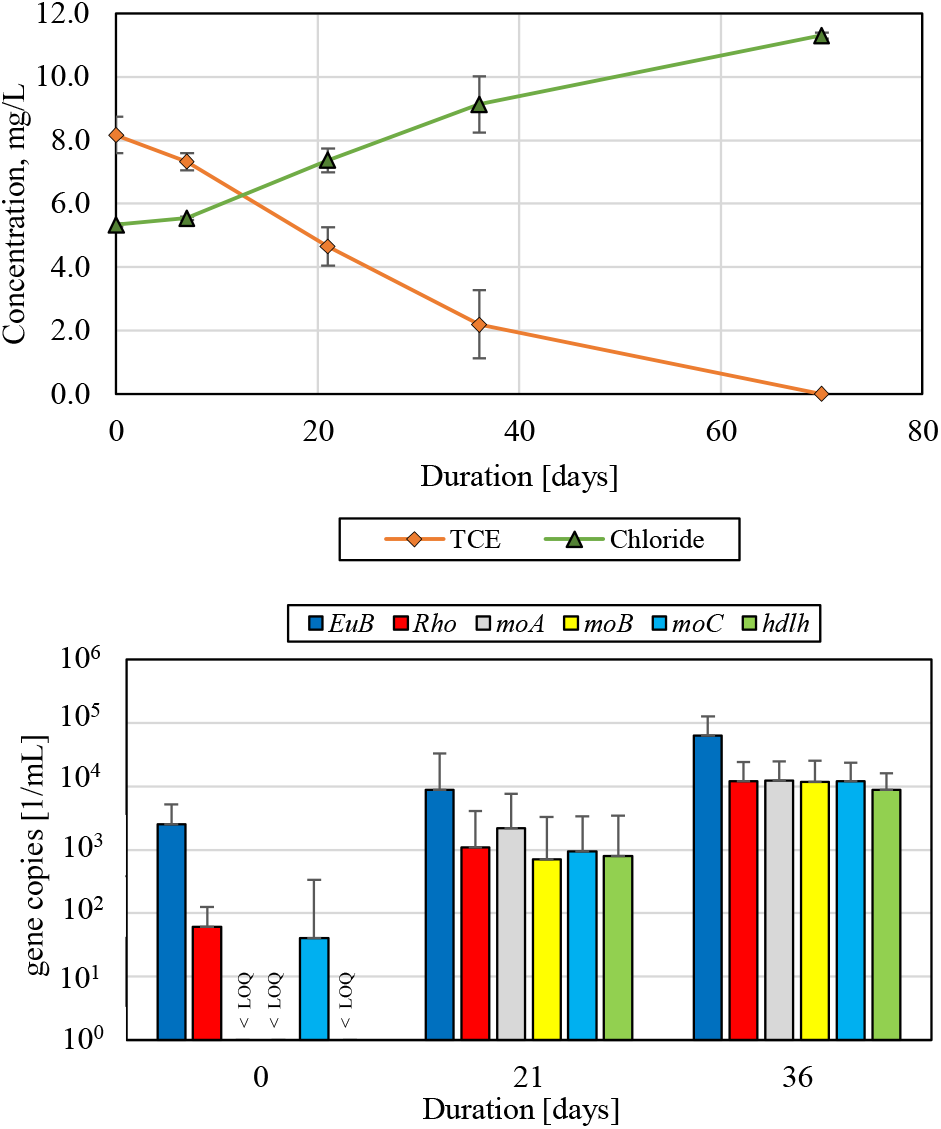
Top: Development of TCE and chloride concentration in the batch experiments. Bottom: Development of gene copy-numbers measured with primers EuB (bacterial 16S rRNA gene), Rho (16S rRNA gene of Rhodocyclaceae), moABC (monooxygenases) and hdlh (haloacid dehalogenase). Error bars indicate standard deviation (σ) of triplicate measurements; “<LOQ” indicates values below the limit of quantification.

The same co-occurrence between TCE degradation and the increase of the gene copies was also observed in second laboratory batch experiment, which was performed in duplicate (Figure S 4).

### 3.4 Primer specificity

To test the specificity of the established primers, different aerobic chloroethene degrading cultures and a wide array of environmental samples have been analyzed. These include samples from municipal wastewater, tar oil contaminated groundwater from different sites containing monoaromatic hydrocarbons (BTEX), polycyclic aromatic hydrocarbons (PAH) and heterocyclic hydrocarbons (NSO-HET), and TCE-contaminated site with an ongoing aerobic bioremediation process. The aerobic metabolic VC-degrading culture was enriched from a (TCE, cis-DCE, and VC) chloroethene-contaminated site in Baden-Wuerttemberg, Germany, and cDCE-degraders were enriched from a (TCE, cis-DCE, and VC) chloroethene-contaminated site in North Rhine-Westphalia, Germany. The *Rhodocyclaceae* (*Rho*) biomarker was quantified in all aerobic chloroethene-degrading enrichment cultures except that of the VC degradation enrichment. It was also quantified in most of the environmental samples without chloroethene pollution. The qPCR analysis targeting the *Rhodocyclaceae* family was expected to be measurable at sites contaminated by other pollutants as *Rhodocyclaceae* are common environmental bacteria. In contrast, the qPCR targeting the functional genes appeared to be highly specific, since they were only detected at sites and in samples with demonstrated aerobic TCE degradation. These results emphasize the suitability of the functional genes as biomarkers for detection of aerobic metabolic TCE degradation potential and monitoring of aerobic remediation processes.

## 4 Conclusion

Amplicon and metagenomic approaches were used to identify the bacterial family of *Rhodocyclaceae* as the putative bacterial taxon responsible for the aerobic metabolic TCE degradation in enrichment cultures and in the groundwater. In addition, the metagenomic bins provided possible insight into the functional genes implicated in the mineralization of TCE. These functional genes encode for monooxygenase and dehalogenase related enzymes. These approaches successfully provided sequences used to develop specific qPCR-methods for the assessment of the aerobic metabolic TCE degradation. The qPCR primers provide a tool for the future investigations of contaminated sites, bioremediation efforts and further research concerning the aerobic metabolic TCE degradation pathway. Attempts at isolating the bacteria responsible for the TCE degradation can now be pursued using the putative genome analysis to determine possible required cofactors derived from other members of the enrichment.

## 5 Author Contribution

A.R.V., A.B., S.H., L.S, and A.T. collected, analyzed, evaluated and interpreted the data, wrote the initial draft and revisions of the manuscript. A.R.V. and J.H. performed the bioinformatics parts of the study. A.B., S.H., and A.T. designed and conducted the experiments. A.T, T.M.V., X.Z. and H.-P.Z. provided resources, acquired funding, reviewed and edited the manuscript. All authors have read and approved the final manuscript.

## 7 Acknowledgements

The authors acknowledge Anna Willmann and Marie Schupfer for their assistance.

## 8 Data availability

Data can be made available on request. Sequences described above have been submitted to public databases with their accession numbers.

## 9 Declarations

Ethics approval: not applicable

Consent to participate: not applicable

Consent for publication: not applicable

Competing interests: The authors declare no competing interests

