## Supplementary material for "Metagenomics and qPCR analysis of aerobic metabolic TCE degrading bacteria": Supplemantary_Information

### S1. Origin of samples for taxonomic and metagenomic analyses

The degradation curves of the batches and groundwater microcosms used in this study are given in Figure S1. The degradation curve of MC-GW can as well be found in the SI material of Willmann *et al.* (2023) (site 5).

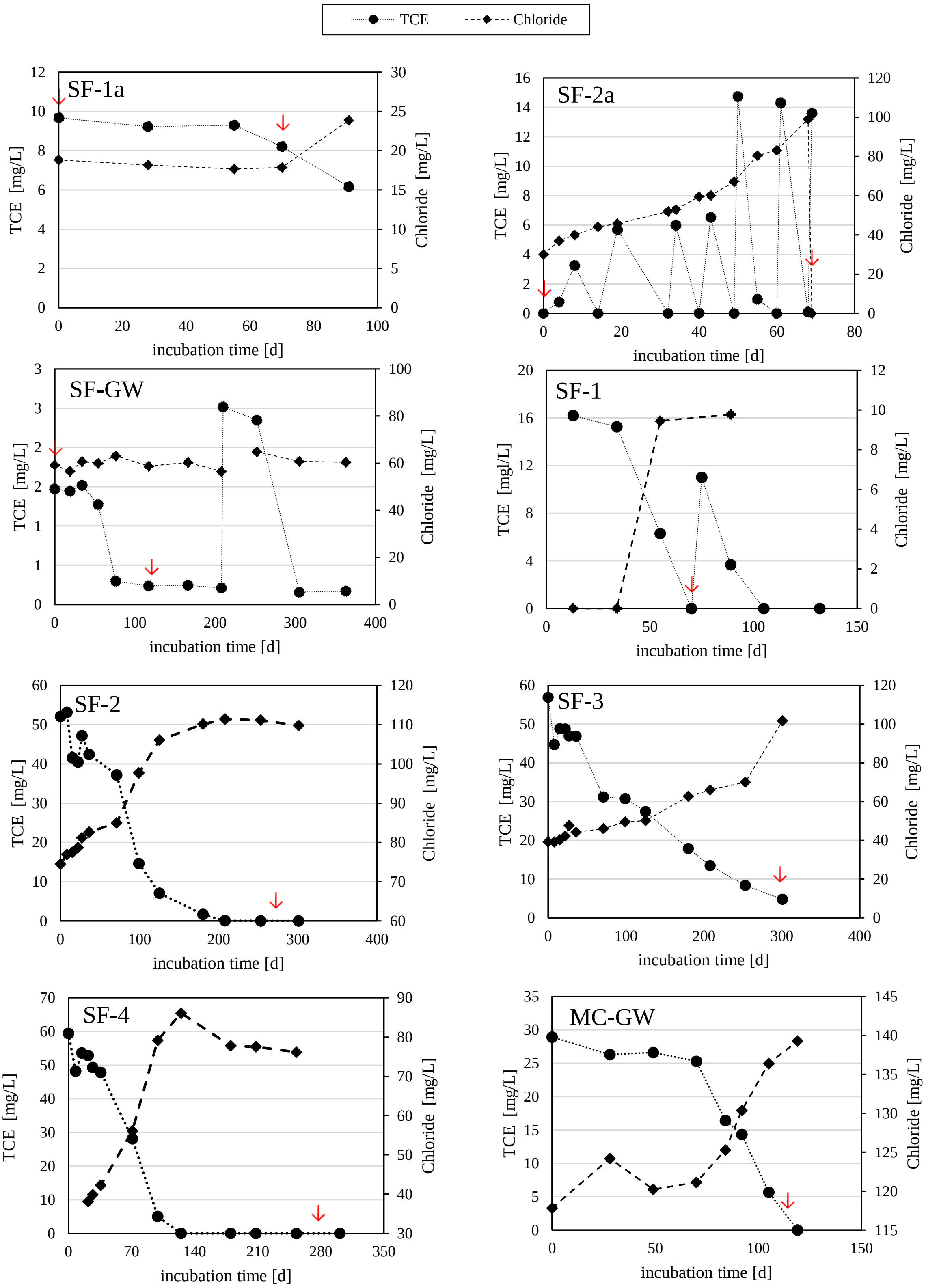


*Figure S1 Degradation of TCE and formation of chloride in the batches used for sequencing analyses. Red arrows indicate the time of sampling for the amplicon and metagenomic analyses.*

**S2. Analysis Bioinformatics workflow**

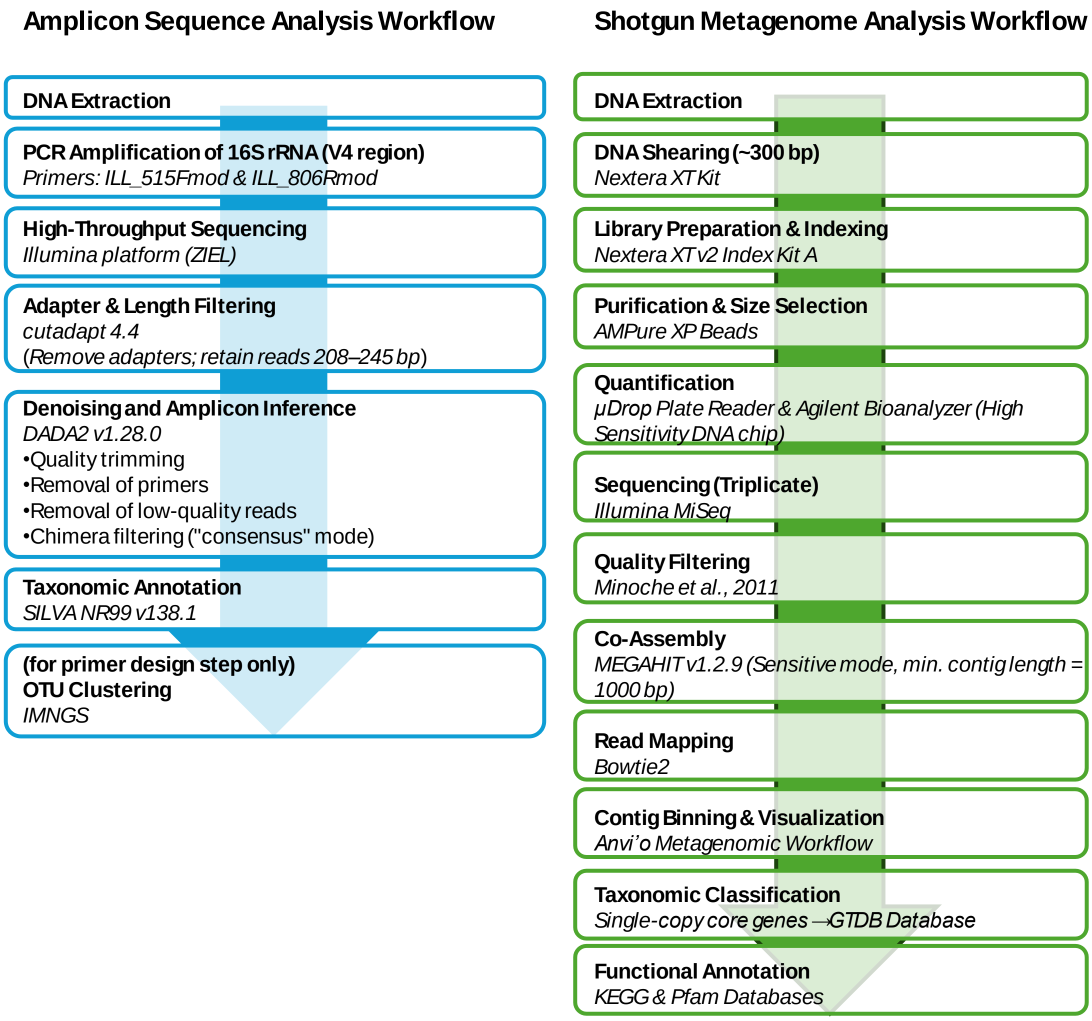


*Figure S2 Bioinformatics workflow including all software tools used and outputs.*

**S3. Database sequences used for the identity of functional genes:**

1: Ammonia monooxygenase subunit A [EC:1,14,18,3 1,14,99,39]

ATGAGTGACGCAAATGCAGTAAATGTAGCTGCGGGGTTGTCGCCAGAAGCGGCCAAAATGGGCCGCTGGATGGATATCATAC
TTATTCTGACAGTTTTGTTCGCGCTTATTGGCGCGTTTAATATTCATTACGCGGTATTCGCAGGAGAGTGGGATTTCTGGGCGG
ACTGGAAGGATCGCGAATACTGGACAGTAGTGACACCAATCGTAGCTATTACCTTTCCGGCCGCAGTACAGAATATTCTGTGG
AGGAACTTCCGTTTGCCTTTCGGTGCCACACTCTGTATCGCTGGACTGCTGCTTGGCGAGTGGCTGGTGCGTATTTTTGGTTTT
GGTATGTGGTCAGGCTATCCGATTAACCTGGTCATGCCGTCACAGCAGATGGGCACTGCGCTTATTCTCGACGCTATCCTCCT
GCTGACGCGTGGCAACTGGATATTCACGGGGGTTTTTGGTGGTGCATTGTATGGTGTACTGTTTTATCCGATGAATTATCCGCA
GATTGCGCCTCTGCATGAGCCGGTAAATTACCAAGGTGTCGTGCTCTCGCTGGCTGATCTCATCGGCTTCATGGATGTCCGTA
CCGGTATGCCAGAGTACATTCGCATGATCGAGCGTGGCACCCTGCGCACGTTTGGTGGTCATTCGACAGCCCTTGCGGCAATT
TTCGCCGGATTCATCTGCATCATGATGTACTGGTTGTGGTGGTACATCGGTGGTCTCTTCTCGAATACGGCTCACATGATCAAA
GGCGTATCAACGCCACAGGTGGATACCGCCGCAGAAGTCGATCACACCACCATTGCAGGAAGGATCTGA

2: Monooxygenase subunit B protein

ATGAAATTCAACATATTCAATAAGCTTAAGAAAGCTGCATTTATTACAGCAACGACTGTAGTCGGGCTACTGTCGTTGGCTGC
GTCTCCTGATGCAATGGCCCATGGCGAACGGGCACAGGAGCCCTTCTTGCGCATGCGCTCGATTCAGTGGTACGACATGCAAT
GGAGTAAGGATACCGTCAAGGTCAATGATGAAATGACGCTGAGTGGGAAGTTCCGTGTTTTCCCCGACTGGCCGAATGCGCT
TCATGAGCCGAAGGTATCTTTTTTGAATATCGGCATTCCTGGCTCGGTATTTTTGCGTACCTCGACTGAAATCGGCGGTGAGGT
ACAGTTTTTCTCCGGCCCGCTGGAAATTGGCAAAGACTACTCTTTCAAGGTCACCGTAAAGGCGCGCAAGCCAGGTGAATACC
ACATGCATTCTCAACTCAACATGTTGAGCAATGGCCCTGTCGCCGGTCCCGGTTCCTGGGTCAAGGTTGTGGGTGACGAGAAG
GATTTCAGCAACCCGATAACGACGCTGACTGGCCGTACCGTCGACACCATGACCGTCGGCAAGGCGGAGGGCATCACATGGC
ATATCATCTGGGCCGTCTTTGGCGCTGCTTGGTTGCTGTACTGGCTCACCCAGCCGCTGTTCCTGGCACGTTTCCGTGCACTGA
ACAACGGACGGGAGGATGTACTGGTAACGTCGGGCCACATGAAGCTCGGTGTGGCGGTATTGGTTGTTAGCCTTGGTTTGATT
TTCTACGGCTATCAGTCGGCCACTGCAAAATACCCGAACGTTGTACCGCTCCAGTCGGGCAAGGCTGTATACAAGTCTATTGA
TCTGCCCCAGTCACCAGCCGTGGTTACAGTTGAAAGCGCGAACTATGATGTACCTGGTCGCGCCATGCGGATTACTGCGACGA
TCACCAACAAGGGTGACAAGCCGTTGCGCATTGGCGAGTTTTCGACAGCCGGTGTGCGCTTCCTCAATCCTGAGTCCGGCATT
CCGCTTGACAATCCGACCTATCCGAAAGAGATGCTTGCTCTGCAAGGCCTGAAGCTGGATAACAATGCTCCGATTCAGCCTGG
CGAAACACGCAAGATGGTCATTAGTGCAGCCGATGCTATCTGGGAACTCCAGCGCTTGGTCGATCTGGTGAATGATCCGGAT
AGCCGGTTTGGTGGCATGCTGTATTTCTATGATGATACTGGCAGGCGTGACATCGTCAGCGTTAGCGGTGCTGTGGTTCCAAT
CTTCACTGCGCTGAAGCTCTGA

3: Ammonia monooxygenase/methane monooxygenase, subunit C

ATGAAACGTGATGCCTTGATTACGAGGGCTACTTGGGCAGTCGAGCAAGGTTCGCTTCTGATTGCTGTGGCGGGTGTTGCCGT
CATGCTCTTGGTTGGCTTGTATCTGGCAGGCTGGATTGGCAGCGAGACAGCCGCAGGTTCGCGGGAATCTTTCTCGCCCGGAA
GCATGTTTGGTTATCTGCCGATATGGCTTGCGGAAGCCATGTTTACGCTGTTTTTATGGCGCCGGATATGGCAGACGCAGAAT
GTGGCGAATATGGCTGCCGGTGGCGACAAGCGCGTAGCGCAAGCCTTTCTTCTCATGATCTGGCTGAGCCTTGATTTACTCAT
GGTTGTTTTTGGGGGCGGCTTGCTTGCGCATATGGATGCTCCTTGGGTACAGATGAGTACTGCGGCTATCAGCCCTGCGCATC
TGGTGACTTTTATTGCCGTAATGCCCGCATATATCGTCTTTGGCGTTTCGTCCTGGCTGTTTGCGCGTACTCGATTGCCTGAGTT
TGCTAAGGGGAATCTCGACCATTTCGTGATTGTCACCATTGCTCCCTTCATGTTTCTGCCAGCTTTCGATCCGCAAGACTTGAT
CCATTCGCTCGATCCAAACAGTAATTTGTACCTGGTTATCTACTGGGTCATAACAATCGCTTGGGTTGTCAATGTTGGTTGGTT
GTTCACGAAACTATTTCGCTATGCGTTGCTCCAATTCCCAAGGAGCGGGTAG

4: Halo acid dehalogenase-like hydrolase

ATGTCTAAGAAGTTCATCCCTATGGCGATTGCCTATGACTTCGACGGGACGCTCGCACCGGGGAATATGCAGGAGTACGACTT
TATCCCCGCCCTAAAGCTCCCCTCAAAACATTTCTGGGACAAGGTTAACGAGCTTGCGAAAAAGCATGAGGCCGACCCGATC
CTTGTGTACATGTATGCCATGCTGGAAGAAGCTCGCACCGCAGGCCTTCCCGTCCGCAAAGCCGACTTCAAGAATTATGGGGT
GAATATCGAACTGTTTCCGGGCGTCAAGGAATGGTTCAAGCGCATCAACGACCATGCCAAGACCAAAGGAATTCGGCTGGAG
CATTTCATCATTTCATCGGGTATCCGCGAGATGGTTGAGGGCACGCCGATTCACAAGGCGTTCAAGAAAATTTATGCGTCGAG
TTTTGTGTTCGACGCAAACGGCGTTGCCTGCTGGCCCGCGCTGGCGATCAACTACACGACCAAAACTCAGTACTTGTTTCGCG
TCAACAAAGGCAGCTTGGATGTTCACGACAATTCCGTGATCAATAAATTTGTGCCGAAGGAGCAACGCCCCGTTCCATTCGAG
CACATGATTTTTATCGGCGACGGCGAAACCGATATTCCATGTATGCGCCTGGTGAAAGATCAGGGTGGGCATTCCATTGCGGT
CTACAACCCCGCCCGGCGAGGCCACAAAAAACATGCCGAGCAATTGGTCAAGGATGGGCGTGCAACATTGGCAGCTACCGCC
GACTATCAGGATGGAGGCCCGATAGATCGGGCTGTAAGAGCCATCATTGACAAGATTGAAGCATCCGCAAGGATTGCCTAA

### S4. Data of functional genes involved in TCE aerobic degradation pathways

The enzymes identified within the sequences / genomes resulting from the metagenomic analysis are shown in Figure S3.

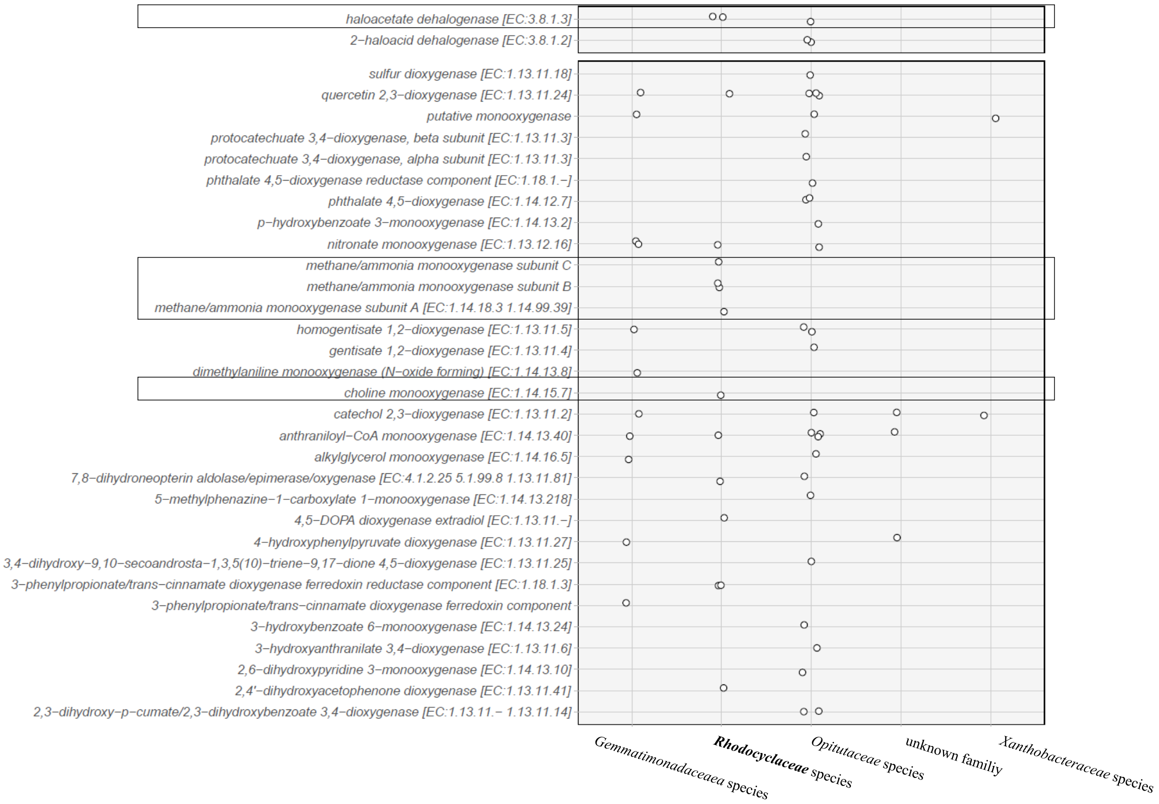


*Figure S3 Occurrence of sequences annotated for mono- and dioxygenases and dehalogenases in the different species. Each dot represents a single gene found in the sequence. The genes coding for the monooxygenases and the dehalogenase are outlined, which were present in the Rhodocyclaceae allocated sequences and were used for specific qPCR development.*

### S5. Correlation of TCE degradation and qPCR analyses – further results

Besides the batch experiment shown in Figure 5, another batch was investigated at seven points with the established qPCR methods and the increase of gene copies could be confirmed as shown in Figure S4.

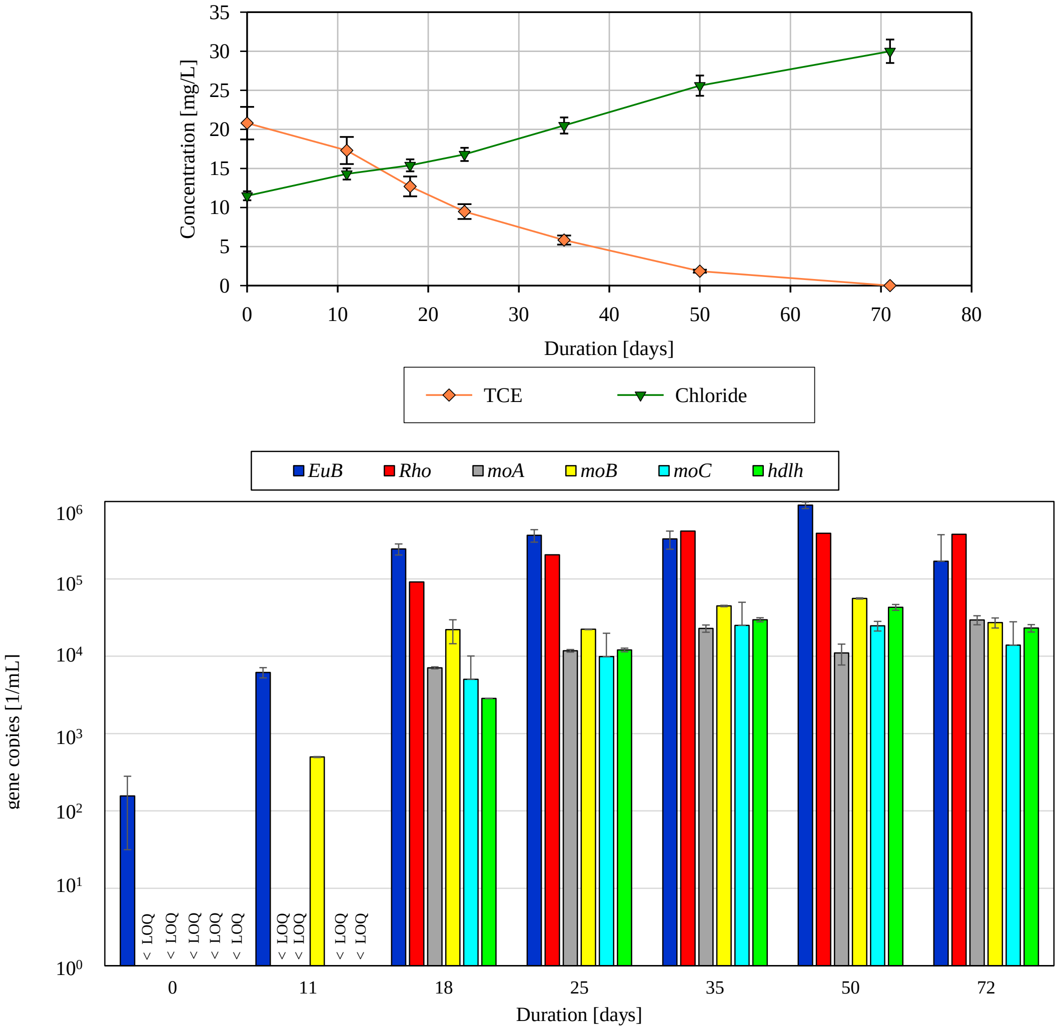


*Figure S4 Top: Development of TCE and Chloride concentration in the batch experiments. Error bars show the standard deviation (σ) of the TCE analysis and standard error (σn) of the Chloride analysis (5%). Bottom: Development of gene copy-numbers measured with primers EuB (bacterial 16S rDNA), Rho (16S rDNA of Rhodocyclaceae), moABC (monooxygenases) and hdlh (halo acid dehalogenase like hydrolase). Error bars indicate standard deviation (σ) of duplicate measurements; "<LOQ" indicates values below the respective limit of quantification.*

### S6. Contaminated site samples used for primer validation

Analytical data are given for the different aerobic chloroethene degrading cultures and the environmental samples (municipal wastewater, tar oil contaminated groundwater from different sites containing monoaromatic hydrocarbons (BTEX), polycyclic aromatic hydrocarbons (PAH) and heterocyclic hydrocarbons (NSO-HET), TCE contaminated site with aerobic bioremediation process).

*Table S1 Analysis data of the samples that were used for testing the specificity of the established qPCR methods. n.a. indicates analyses are not available.*

| Sample | $O_2$, mg/L | pH | $NH_4^+$, mg/L | DOC, mg/L | Sum CE, µg/L | Sum BTEX, µg/L | Sum PAH, µg/L | SUM NSO-HET, µg/L |
|---|---|---|---|---|---|---|---|---|
| Aerobic VC degrading culture (1) | ~8 | 7.2 | n.a. | n.d. | 20,000 | n.a. | n.a. | n.a. |
| Aerobic VC degrading culture (2) | ~8 | 7.2 | n.a. | n.d. | 20,000 | n.a. | n.a. | n.a. |
| Aerobic cDCE degrading culture (1) | ~8 | 7.2 | n.a. | n.d. | 20,000 | n.a. | n.a. | n.a. |
| Aerobic cDCE degrading culture (2) | ~8 | 7.2 | n.a. | n.d. | 20,000 | n.a. | n.a. | n.a. |
| Active TCE degrading site (Sample 1) | 12.72 | 6.9 | <LOQ | 0.87 | 0.88 | n.a. | n.a. | n.a. |
| Active TCE degrading site (Sample 2) | 4.25 | 7.1 | 0.08 | 0.8 | 0.012 | n.a. | n.a. | n.a. |
| Aerobic TCE degrading culture | n.a. | n.a. | n.a. | 4.9 | 24 | n.a. | n.a. | n.a. |
| Municipal waste water | n.a. | n.a. | n.a. | n.a. | n.a. | n.a. | n.a. | n.a. |
| BTEX/ PAH/ NSO-HET contaminated site 1 | 0.9 | 7.0 | 0.82 | 3.6 | n.a. | n.a. | n.a. | n.a. |
| BTEX/ PAH/ NSO-HET contaminated site 2 (Sample 1) | 0.13 | 7.6 | 4.1 | n.a. | 330 | 2770 | 4100 | 1200 |
| BTEX/ PAH/ NSO-HET contaminated site 2 (Sample 2) | 0.09 | 7.5 | 0.97 | n.a. | 1.0 | 0.4 | 21 | 4.8 |
| BTEX/ PAH/ NSO-HET contaminated site 3 (Sample 1) | < 0.5 | 7.7 | 5.4 | 10 | n.a. | 960 | 11,000 | 2,200 |
| BTEX/ PAH/ NSO-HET contaminated site 3 (Sample 2) | 0.8 | 7.1 | 8.1 | 2.5 | n.a. | 13 | 300 | 32 |
| BTEX/ PAH/ NSO-HET contaminated site 3 (Sample 3) | 2.4 | 6.5 | 0.73 | 1.3 | n.a. | <LOQ | 0.015 | 0.024 |
